# Between two walls: Modeling the adsorption behavior of β-glucosidase A on bare and SAM-functionalised gold surfaces

**DOI:** 10.1101/2021.07.02.450859

**Authors:** Nicolas Bourassin, Florent Barbault, Marc Baaden, Sophie Sacquin-Mora

## Abstract

The efficient immobilization of enzymes on surfaces remains a complex but central issue in the biomaterials field, which requires us to understand this process at the atomic level. Using a multi-scale approach combining all-atom molecular dynamics and coarse-grain Brownian dynamics simulations, we investigated the adsorption behavior of β-glucosidase A (βGA) on bare and SAM-functionalized gold surfaces. We monitored the enzyme position and orientation during the MD trajectories, and measured the contacts it forms with both surfaces. While the adsorption process has little impact on the protein conformation, it can nonetheless perturb its mechanical properties and catalytic activity. Our results show that compared to the SAM-functionalized surface, the adsorption of βGA on bare gold is more stable, but also less specific, and more likely to disrupt the enzyme’s function. This observation emphasizes the fact that the structural organization of proteins at the solid interface is a keypoint when designing devices based on enzyme immobilization, as one must find an acceptable stability-activity trade-off.

**TOC image:** 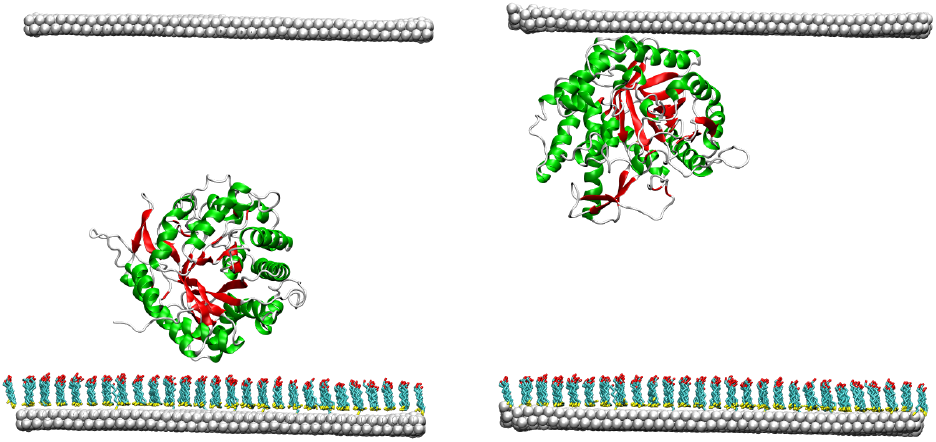

## 1. Introduction

The immobilization of enzymes on solid supports has attracted growing interest in the biomaterials field over the last forty years,^1–2^ as this phenomenon plays a central part in numerous applications,^3–4^ such as biosensors,^5–6^ biofuel cells^7–9^ or biomedical devices^10–11^. When investigating the interaction between an enzyme and a solid surface, two key issues that should be addressed are the stability of the protein/surface interface, and the protein orientation on this surface. Ensuring a correct protein orientation is essential for example in bioelectrocatalysis devices, where it will enable a direct electron transfer between the adsorbed redox enzyme and the electrode.^12–15^ In addition, one must make sure that the enzyme active site remains accessible after immobilization.^16^ This can be achieved either through targeted chemical linkage between the protein and the surface,^17–19^ or via the adequate functionalization of the surface, for exemple with self-assembled monolayer (SAMs), in order to adapt its charge and physico-chemical properties to a specific protein.^20–23^ One must also pay attention to the conservation of the adsorbed protein’s structure and dynamics, as perturbations in the protein conformation^24–26^ or internal mobility^27^ are likely to result in a dramatic decrease of the catalytic activity. Altogether, the molecular mechanisms taking place upon enzyme immobilization are not well understood. While most experiments on immobilized proteins, either via a covalent^28–29^ or non-covalent^30–32^ strategy, report a decrease in their catalytic activity, sometimes enzyme immobilization can also result in an important increase of the catalytic performance.^33^

Numerous enzyme immobilization strategies have been developed, that involve simple adsorption via electrostatic or van der Waals interactions, the use of porous materials, crosslinking, or multipoint-covalent attachment,^33–35^ and many experimental approaches are now available to investigate protein immobilization on solid surfaces, including atomic force microscopy, mass spectrometry and spectroscopic methods.^36–37^ Still, experimental techniques fail to provide us structural information regarding the protein-surface interaction at the atomistic level. As a consequence, computational models have been playing an increasingly important part, as they can bring greater hindsight on the chemical and biological processes taking place at the bio-nano interface. The last decade has seen a wealth of methodological developments, in particular with multiscale approaches combining all-atom and coarse-grained representations.^38–41^ Molecular simulations techniques are now a powerful tool in the biomaterials field, ^42–45^ as they enable us to model various surface types^22, 46^ (such as metals, oxides, carbon or silica-based of functionalized with SAMs), and investigate how the surface structure^47^ and chemical properties can impact the orientation, conformation and activity of immobilized proteins.

β-Glucosidase A (βGA) is an enzyme which cleaves β-glucosidic linkages in disaccharide or glucose-substituted molecules, and plays a fundamental role in processes such as cellulose degradation.^48^ The enzyme is extensively used by the industry for the production of ethanol for biofuel, and the optimization of its function and reusability is thus of practical concern.^49–53^ Experimental studies on immobilized βGA, using covalent^28, 50^ or non-covalent^51–52^ binding strategies, show that the immobilized enzyme presents an enhanced thermal stability and pH tolerance, but a reduced catalytic activity compared to the free enzyme. This decrease in the enzymatic function after immobilization can be due to several factors, such as a reduced accessibility of the substrate to the catalytic active site, or changes in the enzyme conformation and rigidity.

In this work, we combine all-atom Molecular Dynamics (MD) and coarse-grain Brownian Dynamics (BD) simulations to investigate the adsorption behavior of βGA, namely its orientation with regard to the solid surfaces and the impact of adsorption on its structure and dynamics, when it is confined between a bare and a hexanol-thiol-SAM-functionalized gold surface. While the analysis of the MD trajectories shows that βGA will preferentially adsorb on the bare gold surface, the coarse-grain calculations suggest that this stronger protein-surface interaction is also more likely to perturb the enzyme mechanical properties, which in turn might have some repercussion on its function. Altogether, our results highlight the fact that when setting up a protein immobilization strategy, one must achieve a delicate balance between the enzyme stability and its catalytic activity.

## 2. Material and Methods

### All-atom Molecular Dynamics simulations

#### Simulation setup

The 3D coordinates of the βGA were extracted from the protein data bank under the accession code 3ahx, which corresponds to the X-ray structure of bacterial βGA from *Clostridium cellulovorans*.^48^ Molecular dynamics simulations (MD) for the protein/surface system were realized with the Amber 16^54–55^ software suite under periodic boundary conditions with the Particle Mesh Ewald (PME) method.^56^ The enzyme’s net charge is −13e, and 13 Na^+^ ions were randomly placed in the simulation box to neutralize the system. We then used the center of mass of the βGA heavy atoms as the center of the referential for calculating the enzyme’s dipole moment,^57^ with the x and y axes being parallel to the solid surfaces, and the z axis perpendicular to it (see Figure 1). βGA has a weak dipole moment (of around 20D) and its electrostatic surface (shown in Figure SI-1) does not present any large positively or negatively charged patch. As a consequence, and unlike proteins with a large dipole moment such as hydrogenases^12^ or bilirubin oxidases^58^, it is not possible to infer its orientation on a polar surface from its electrostatic properties. To explore how the starting protein orientation will impact its adsorption on the solid surfaces, six different orientations were generated like a dice, by rotating the protein around the x and z vectors (shown in Figure 1), so that each starting orientation shows a different face of the enzyme toward the functionalized surface, and the enzyme is initially placed in the simulation box with its center of mass 3.2 nm above the SAM-functionalized surface. The six starting orientations have been labelled as z0, z90, x0, x90, x180 and x270 (and are shown in Figure SI-2), and the corresponding simulations will be referred to as the *confined trajectories* for the rest of the manuscript, as the displacement of the enzyme in the z-direction is limited by the solid surfaces.

**Figure 1:**
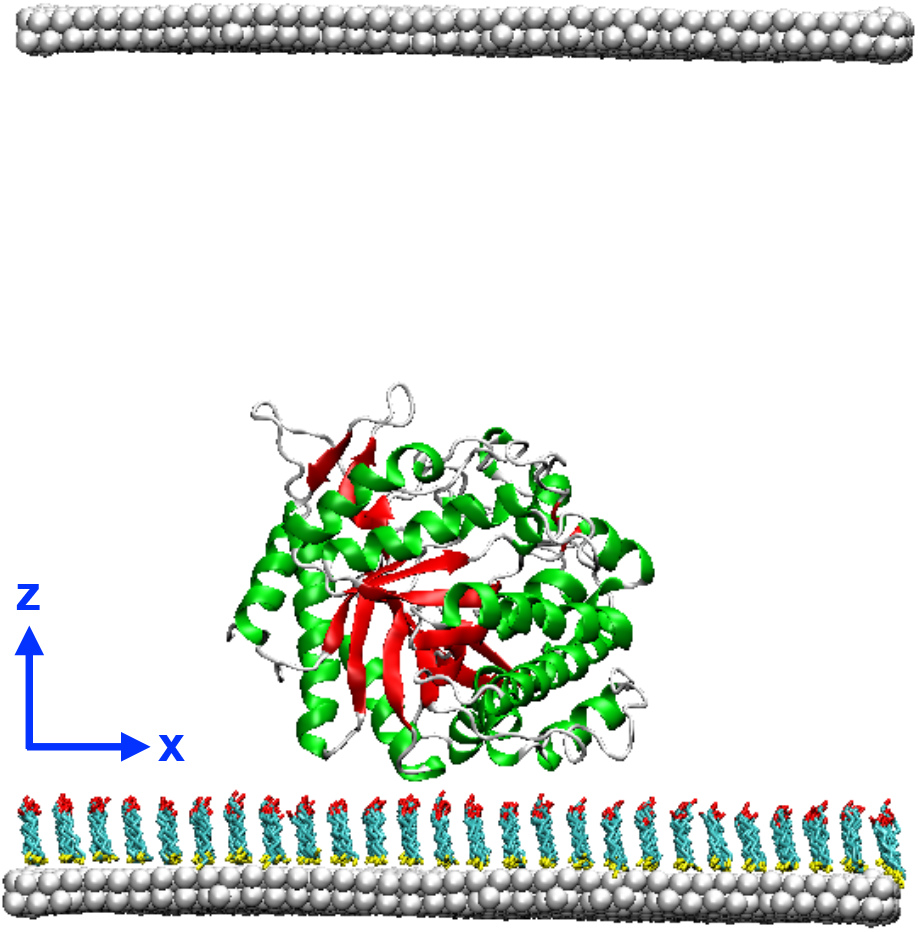
Initial position and orientation of βGA in the z0 trajectory. The five additional starting orientations were obtained by rotating the enzyme around the x and z axes. Figures 1, 4 and 6a (and SI-1, SI-2, SI-3 and SI-5d) were prepared using Visual Molecular Dynamics.

Consideration of the gold surface functionalized with a 6-mercapto-1-hexanol SAM were reproduced from previous works.^59–60^ Starting from this last validated model, the surface was extended to a square of 132 nm^2^ in order to let enough room for the protein to make translations and rotations during simulations. To facilitate adsorption analyses, the functionalized gold surface is placed on the xOy plane and periodic conditions are set to provide a uniform surface on the xOy plane. Atomic distances between *real and image space* is of 12.5 nm along the z direction so that the protein is placed in a rectangular simulation box, between two homogenous solid surfaces: a SAM-OH-functionalized one *below* and a solely Au{111} surface at 12 nm *above*. The six starting systems were then solvated with TIP3P water molecules.^61^ The ff14SB force-field^62^ was chosen for the protein while the functionalized gold surface was treated with a previously validated set of parameters.^60^ A technical note on this parameters choice is available as supplementary information. The final system comprises around 7 000 atoms for the enzyme, 16 500 atoms for the SAM-functionalized gold surface and 45 000 water molecules.

The six systems were subjected to the same simulation protocol. First, a two-step minimization of the solvent has been carried out with the surface and the alpha carbons of the protein restrained by a harmonic potential of 10 and 1 kcal.mol^−1^.Å^−2^ respectively. Then, a final optimization without restraints was performed until a gradient convergence lower than 10^−5^ kcal. mol^−1^.Å^−1^ was achieved. The systems were then heated to 300K with a first 100 ps step in the NVT ensemble to 100K, followed by 100 ps in the NTP ensemble to reach 300 K. In order to avoid premature surface adsorption during this heating phase, 10 kcal.mol^−1^.Å^−2^ harmonic restraints on the solute (surface and protein alpha carbons) were added to the systems leading, sometimes, to some undesired vacuum bubbles. To remove them, a supplemental 500 ps NTP equilibration has been performed at 300 K where the surface and protein were kept frozen. During this stage the water density reaches an equilibrated value of 1.07 by adjusting its periodic volume in the z-direction so that the distance between the surface in the real and image spaces is around 12.0 nm.

The six confined MD simulations were performed for 100 ns each in the NVT ensemble, which is classically chosen for solid interface investigations.^32, 58, 63^ During the MD production, positions of Au, S and its first covalent carbon atom were kept restrained (with a harmonic restraint of 10 kcal.mol^−1^) so that distance and tilt angle, determined previously at the QM level,^59^ were respected in the MD simulations. The heating, equilibration and production parts were performed with 2 fs integration steps. 1000 frames (corresponding to one frame every 0.1 ns) were saved for each of the six confined trajectories.

In addition, a MD simulation of βGA in solution was performed with AMBER using a similar protocol. A two-steps minimization was undertaken with a first optimization of the solvent while the solute is restrained with harmonic restraints of 10 kcal.mol^−1^. These restraints are removed for the second minimization until energy convergence. A two-step heating followed, with first 50 ps in the NVT ensemble to reach 100K, and then 100 ps in the NPT ensemble to reach 300K. This last stage was followed by a solvent density equilibration step of 500 ps in the NTP ensemble at 300 K. Finally, a production run of 485 ns was performed in the NPT ensemble. The properties of βGA during this simulation, which we will refer to as the *bulk trajectory*, were used as a reference when assessing the impact of surface adsorption on the protein. Again, we used 2 fs integration steps, and 1000 frames (corresponding this time to one frame every 0.485 ns) were saved for the bulk trajectory.

#### Analysis

The gmx_cluster algorithm from the Gromacs (2018.8) suite^64^ was used to extract statistically representative structures for each trajectory over the simulation production period. The clustering cutoff was modulated in order to obtain between one and three clusters for each trajectory (depending on the amplitude of the conformational changes undergone by the enzyme during the MD simulation), and for each cluster, the most representative frame was kept. Each frame represents a cluster that must comprise at least ten members (1% of the total trajectory). And for a given trajectory, the ensemble of the selected structures must represent at least 90% of this trajectory. Alltogether, for the seven trajectories (bulk and confined) these criterions lead to the selection of 14 frames that are listed in Table1. All the structures remain very close to the experimental reference structure, with a backbone rmsd comprised between 1 and 2 Å. These structures (shown in Figure SI-3) were then used to investigate the mechanical changes undergone by βGA during the simulations.

**Table 1:** Representative structures from the bulk and confined MD trajectories that were selected for mechanical investigation after clustering.

| Trajectory | Clustering cutoff (Å) | Position of the selected frame (1000 frames trajectory) | % of trajectory representation |
| --- | --- | --- | --- |
| **Bulk** | 0.85 | 360 | 99.7 % |
| **z0** | 0.73 | 118 | 19.8 % |
|  |  | 461 | 43.4 % |
|  |  | 814 | 29.8 % |
| **z180** | 0.73 | 19 | 2.4 % |
|  |  | 186 | 93.0 % |
|  |  | 990 | 1.8 % |
| **x0** | 0.73 | 270 | 97.6 % |
| **x90** | 0.73 | 265 | 61.6 % |
|  |  | 913 | 37.9 % |
| **x180** | 0.73 | 62 | 7.4 % |
|  |  | 171 | 34.7 % |
|  |  | 797 | 54.9 % |
| **x270** | 0.73 | 771 | 97.3 % |

### Coarse-grain Brownian Dynamics simulations

#### Rigidity profile of a protein

Coarse-grained Brownian Dynamics (BD) simulations were run using the ProPHet (Probing Protein Heterogeneity, available online at https://bioserv.rpbs.univ-paris-diderot.fr/services/ProPHet/) program^65–67^. In this approach, the protein is represented using an elastic network model (ENM). Unlike most common coarse-grained models where each residue is described by a single pseudoatom,^68^ ProPHet uses a more detailed representation^69^ that involves up to 3 pseudoatoms per residue and enables different amino acids to be distinguished. Pseudoatoms closer than the cutoff parameter *R_c_* = 9 Å are joined by Gaussian springs which all have identical spring constants of γ_struct_ = 0.42 N.m^−1^ (0.6 kcal.mol^−1^·Å^−2^). The springs are taken to be relaxed for the initial conformation of the protein. The simulations use an implicit solvent representation via the diffusion and random displacement terms in the equation of motion,^70^ and hydrodynamic interactions are included through the diffusion tensor.^71^

Mechanical properties are obtained from 200,000 BD steps at an interval of 10 fs and a temperature of 300 K. The simulations lead to deformations of roughly 1.5 Å root-mean-square deviation with respect to the protein starting conformation (which - by construction - corresponds to the system’s equilibrium state). The trajectories are analyzed in terms of the fluctuations of the mean distance between each pseudoatom belonging to a given amino acid and the pseudoatoms belonging to the remaining residues of the protein. The inverse of these fluctuations yields an effective force constant *k_i_* describing the ease of moving a pseudoatom with respect to the overall protein structure.

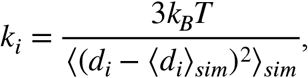

where 〈〉_*sim*_ denotes an average taken over the whole simulation and *d_i_* = 〈*d_ij_*〉_*j*\*_ is the average distance from particle *i* to the other particles *j* in the protein (the sum over *j\** implies the exclusion of the pseudoatoms belonging to residue *i*). The distance between the C_α_ pseudoatom of residue *i* and the C_α_ pseudoatoms of the adjacent residues *i-1* and *i+1* are excluded since the corresponding distances are virtually constant. The force constant for each residue *k* is the average of the force constants for all its constituent pseudo atoms *i*. We will use the term *rigidity profile* to describe the ordered set of force constants for all the residues of the protein.

## 3. Results and discussion

### β-Glucosidase position in the simulation cell

The distance between the enzyme center of mass and the SAM-OH surface during the confined trajectories is shown in Figure 2. While all confined trajectories start with βGA positioned close to the SAM functionalized surface, the enzyme remains adsorbed on the SAM in only two cases out of six (trajectories x0 and z0). For the remaining trajectories, βGA moves away from the lower, SAM-OH covered, surface within the first half (50 ns) of the simulation, and will either remain in an unbound state (traj. x270) or eventually bind to the bare gold surface (traj. z180, x90 and x180). At this 100 ns timescale, the enzyme adsorption on bare gold appears to be significantly more stable than on the SAM-OH surface, as no detaching event was observed in any of these three simulations for the remaining time of the trajectory. This observation can be related to the interaction energies between the enzyme and the two solid surfaces which are shown on Figure SI-4. The energy values were obtained with the NAMDEnergy plugin^72^ and correspond to non-bonded van der Waals and electrostatic terms. As the gold atoms are not charged,^59–60^ the interaction energy between βGA and the bare gold surface is entirely due to the van der Waals energy term. For the polar SAM-OH surface, van der Waals interactions account for roughly 10% of the interaction energy, while the rest is due to the electrostatic interaction term. The observed energy values are significantly more favorable for the protein-bare gold interaction (down to −400 kcal.mol^−1^) as compared to the protein-SAM-OH interaction (down to −150 kcal.mol^−1^), thus accounting for the increased stability of βGA adsorbed on gold. Interestingly, these values are agreement with those obtained in earlier simulation work on proteins adsorbed on SAMs^32, 58, 73^ or bare-gold surfaces.^63, 74–76^ Some studies using different force fields such as CHARMM^77–78^ and OPLS^79–80^ for the proteins and SAMS, or GolP^81^ and coarse-grain models^82–83^ for the gold atoms, thus showing the robustness of molecular simulations approaches for the investigation of protein-surface interactions. These results also concur with earlier experimental work showing the preferential adsorption of proteins on apolar (here the bare gold) substrates compared to polar (SAM-OH covered gold) substrates.^84^

**Figure 2:**
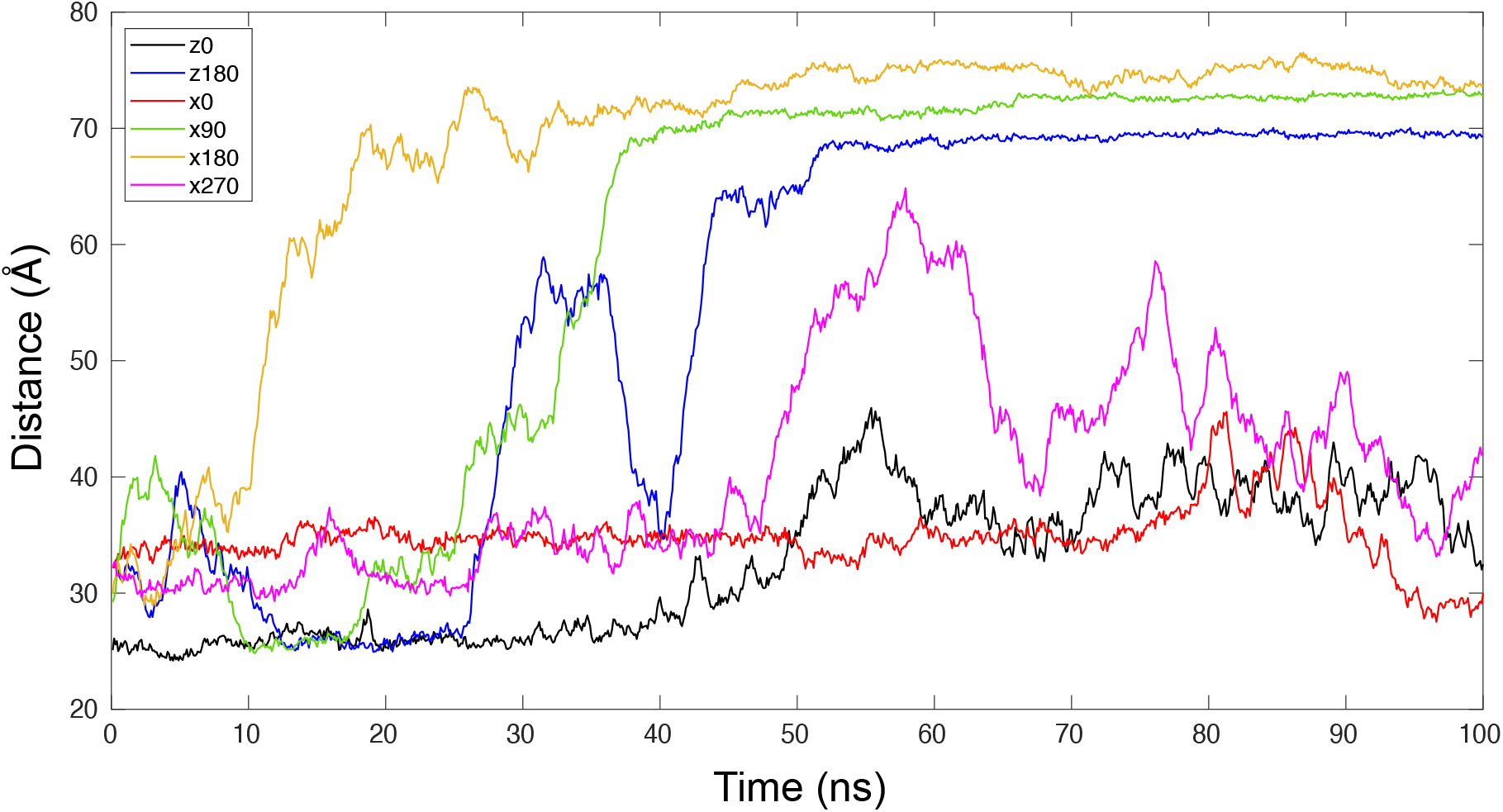
Distance between the SAM surface and the βGA center of mass as a function of time for the six confined MD simulations.

### β-Glucosidase orientation and contacts with the surfaces

βGA presents a small dipole moment (roughly 20D), and its norm and orientation within the protein structure are very stable during all the simulations that we carried out (see Figure SI-5 for the distribution of the dipole moment orientation with regard to the βGA structure along time). Therefore we could use the tilt angle *θ* formed by the dipole moment orientation vector with regard to the SAM-covered surface plane to monitor the enzyme’s orientation. In addition, the enzyme’s rotation around its dipole moment (twist angle, *φ*) was measured by following the vector formed by the Cα atoms of residues Ile119 and Asn353, as it remains perpendicular to the dipole moment over time (see Figure SI-5c). The distribution of the tilt and twist angles along time for the six confined trajectories is shown in Figure 3. All trajectories present a different initial (*φ*, *θ*) orientation. Trajectories z180, x90 and x180, where the enzyme binds to the bare gold surface, cover different parts of the orientational space. On the other hand, trajectories z0, x0 and x270, where the enzyme only binds to the SAM-functionalized surface, seem to converge toward the same area of the graph (highlighted by a black circle in Figure 3). This observation suggests a preferential orientation of βGA on the SAM covered surface, while the stronger (in terms of interaction energy) binding of the enzyme of the bare gold surface is also less specific orientationwise. Interestingly, the bare gold surface bears no atomic charge and makes favorable interactions only through van der Waals attraction, while the SAM covered surface, with hydroxyl moieties on the head, displays a slightly negative surface.

**Figure 3:**
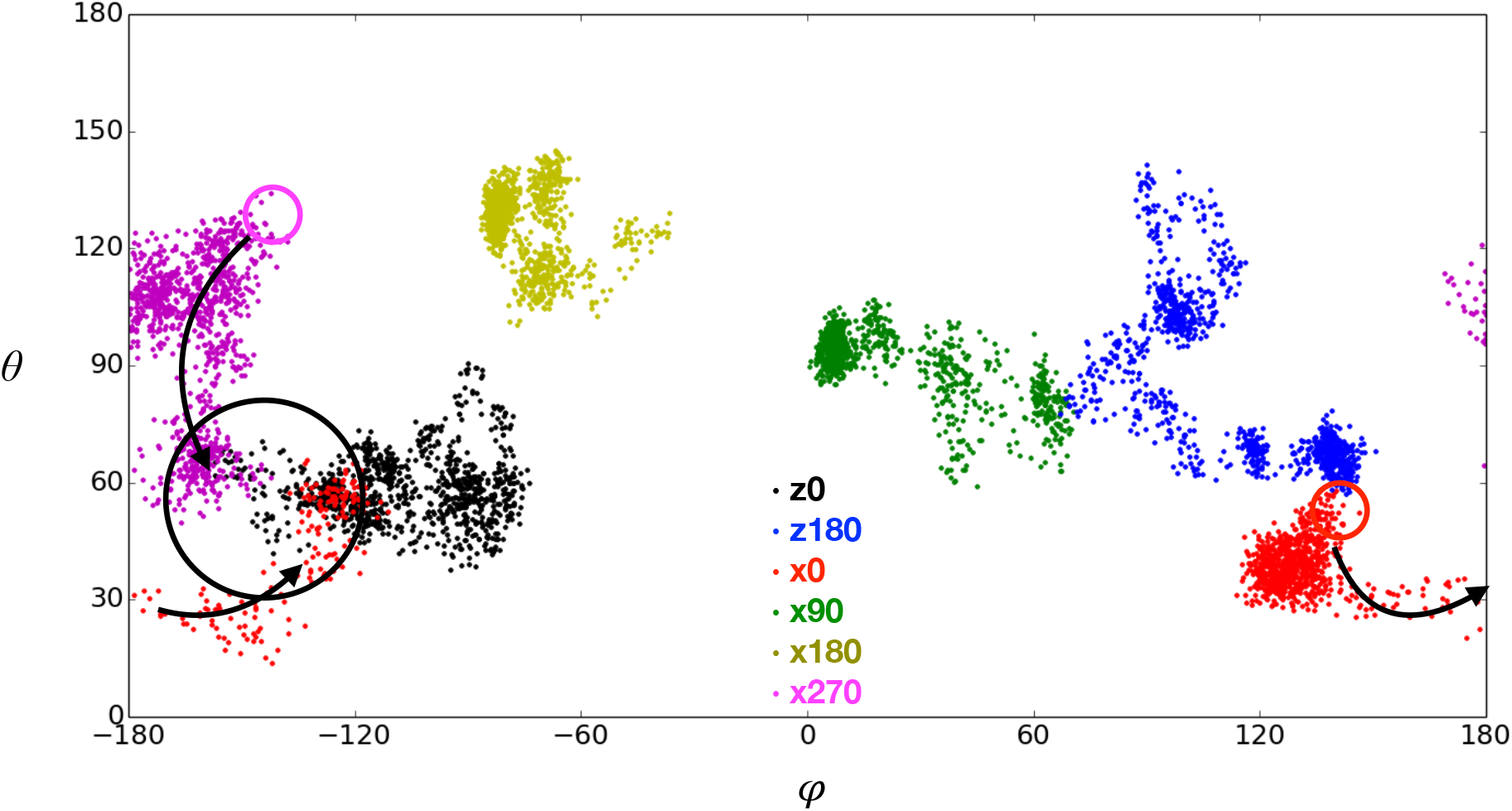
Distribution of the twist (φ) and tilt (θ) angles during the six confined MD simulations. The color code is the same as in figure 2. The red and magenta circles indicate the initial dipole moment orientations for trajectories x0 and x270 respectively, and the black arrows highlight the evolution of the dipole moment orientation along time for these two trajectories.

We investigated this issue further by looking at the parts of the enzyme’s surface forming contacts with the solid surfaces. In our analysis, βGA residues are considered to be in contact with the surfaces when the distance between one of their heavy atoms and a surface atom (SAM or gold) is less than 5 Å. The cumulated contact frequencies for the six trajectories were mapped on the protein surface and are shown in Figure 4 (and the average contact frequencies as a function of the residue number are available in Figure SI-6). βGA binds both solid surfaces preferentially through residues Ala231, Asp232 and Glu233 (shown as a blue patch in Figure 4), which are located on the tip of an α-helix protruding out of the enzyme globular shape. Note that this anchoring point corresponds to one of the most flexible parts of the protein (see Figure 6b), thus allowing the protein to present various orientations, even when it is surface bound. In addition, we can see in Figures 4c and d that the contacts formed between βGA and the bare gold layer seem to be less specific than with the SAM covered gold, as the binding patches are broader, and more widely distributed over the protein surface. In particular, one can observe an additional binding site on the opposite side of the enzyme that seems to be specific to the protein-bare gold interaction.

**Figure 4:**
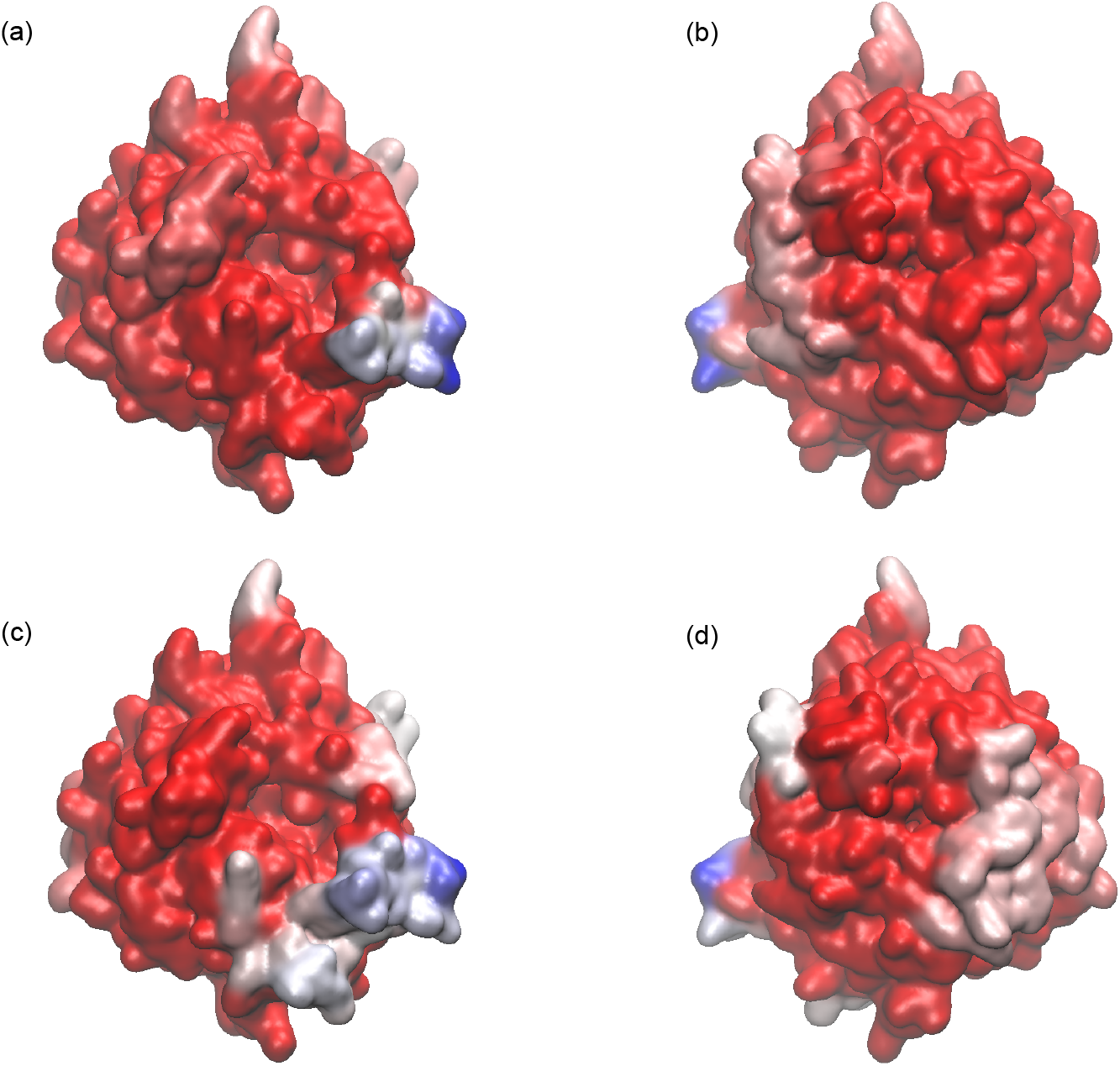
Mapping the contact frequencies between βGA and the gold surfaces on the protein surface; red: low frequency areas, blue: high frequency areas; two views with a rotation around the vertical axis. (a) (b) Contacts between βGA and the SAM-covered gold surface, (c) (d) Contacts between βGA and the bare gold surface.

### Conformational stability

The enzyme overall structure remains stable during all the trajectories, both for the bulk and confined simulations, with a backbone RMSD comprised between 1 and 2 Å (see Figures 5a and b). Besides, the protein central β-barrel structure is strongly conserved, and most conformational changes occur within the enzyme’s outer shell, as can be seen in Figures 5c-f. Only two confined trajectories lead to a noticeable increase of the protein RMSD with regard to the bulk simulation, namely x90 and x180 (which are respectively shown in green and orange in Figure 5). In both cases βGA adsorbs on the bare gold surface. For the x90 trajectory, the interaction with the bare gold surface is particularly strong, as it will even lead to a slight deformation of the protein core (see Figure 5d). The larger RMSD values obtained for the bare gold-bound enzyme compared to the SAM-bound enzyme are in agreement with earlier results obtained by Peng *et al*.^63^ when modeling the adsorption of cytochrome *c* on similar surfaces. In addition, we calculated the enzyme radius of gyration (Rg), which is also very stable, as it remains in the [21.0-21.6] Å range for all simulations. The density distributions (shown in Figure SI-7) are slightly broader for trajectories x90 and x180 (where the enzyme binds to the bare gold surface), and the protein deformations seem to lead to a more compact state, as the Rg distribution is shifted toward smaller values. Inversely, trajectories where βGA remains adsorbed on the SAM layer (z0 and x0) or ends up in an unbound state (x270) present Rg distributions closer to the one obtained for the bulk simulation.

**Figure 5:**
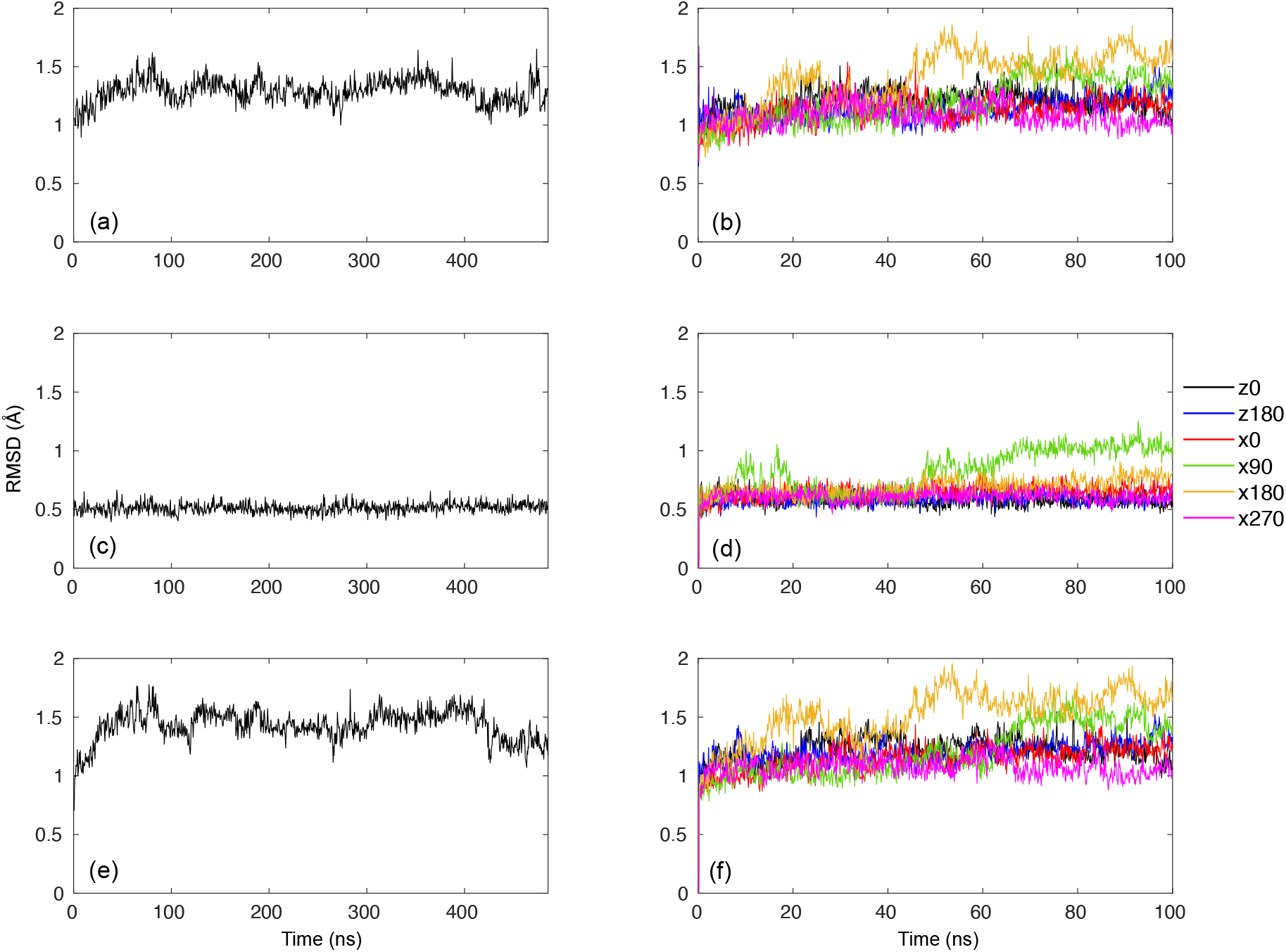
Backbone RMSDs as a function of time, for the whole protein structure (a) βGA in solution (b) confined βGA trajectories; for the central β-barrel only (c) βGA in solution (d) confined βGA trajectories; for the outer shell only (e) βGA in solution (f) confined βGA trajectories.

### Mechanical properties of the β-Glucosidase

#### In the bulk trajectory

The enzyme has a TIM-barrel fold, with a central β-barrel (shown in red in Figure 6a) surrounded by α helices (shown in green in Figure 6a). Its active site is located at the β-barrel entrance and comprises two catalytic glutamate residues, Glu166 and Glu352^48^ (shown as purple spheres in Figure 6a). The rigidity profile obtained for the bulk simulations is shown in Figure 6b and reflects the protein regular structure, with a periodic array of rigidity peaks corresponding to each one of the central β-sheets that is characteristic of the TIM-barrel fold.^66^ The peaks are separated by areas of low, oscillating force constants, that correspond to the external α helical residues. Both catalytic glutamates belong to highly rigid areas in the protein, a classic feature of catalytic residues in enzymes.^85–86^

**Figure 6:**
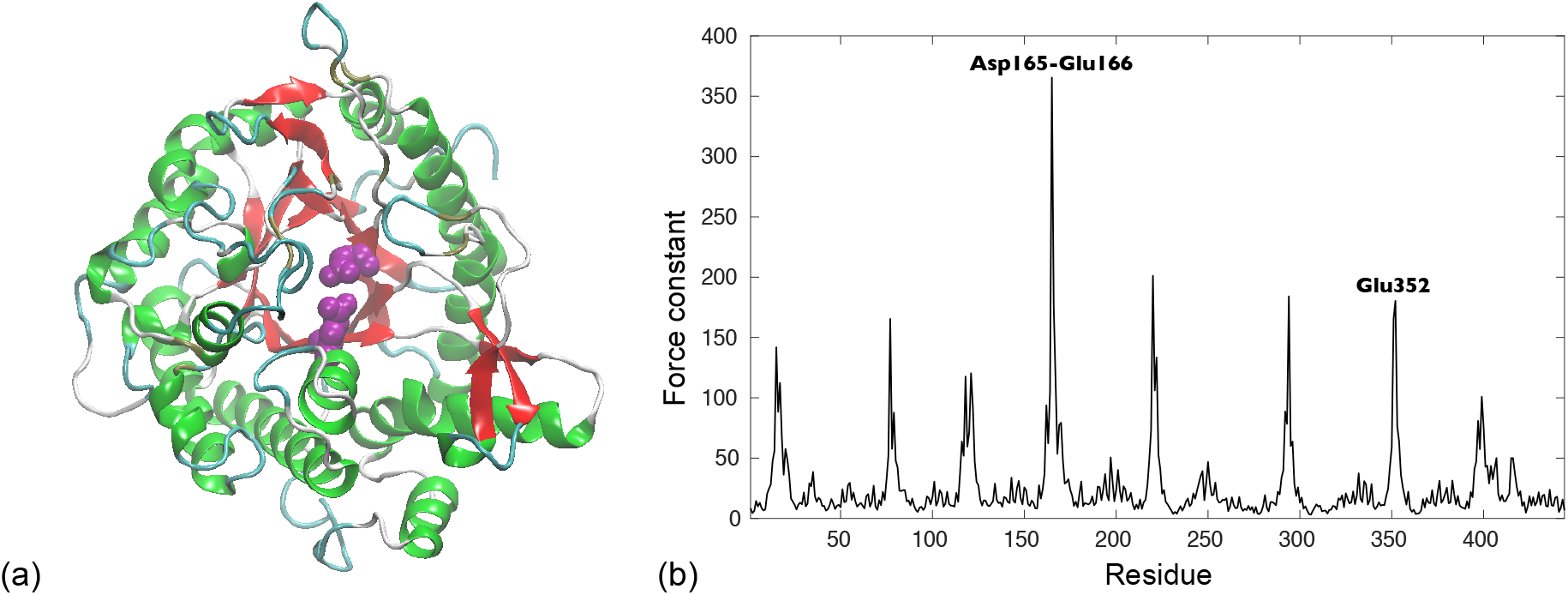
(a) Cartoon representation of βGA, with the central β-barrel shown in red and the external a helices in green. The two catalytic glutamates Glu166 and Glu352 are shown as purple van der Waals spheres. (b) Rigidity profile (in kcal.mol^−1^.Å^−2^) for the βGA in solution.

#### Mechanical variations in the adsorbed β-Glucosidase

We systematically compared the force constants obtained for the 13 βGA representative structures from the confined trajectories and the force constants from the bulk trajectory. The resulting changes in the protein rigidity are shown in Figures 7 (subset of four structures) and SI-8 (all 13 structures). We observe important variations in βGA rigidity that are heterogeneously distributed along the protein sequence. The residues displaying the most important mechanical changes (which can be an increase or a decrease in their force constant) are originally rigid residues from the central β-sheets. This observation concerns in particular Glu166 and Glu352, and two other residues (Asn294 and Tyr296) that also belong to the catalytic site. One should note that these important mechanical variations take place even though the central β-barrel only undergoes minor conformational changes during the simulations.

**Figure 7:**
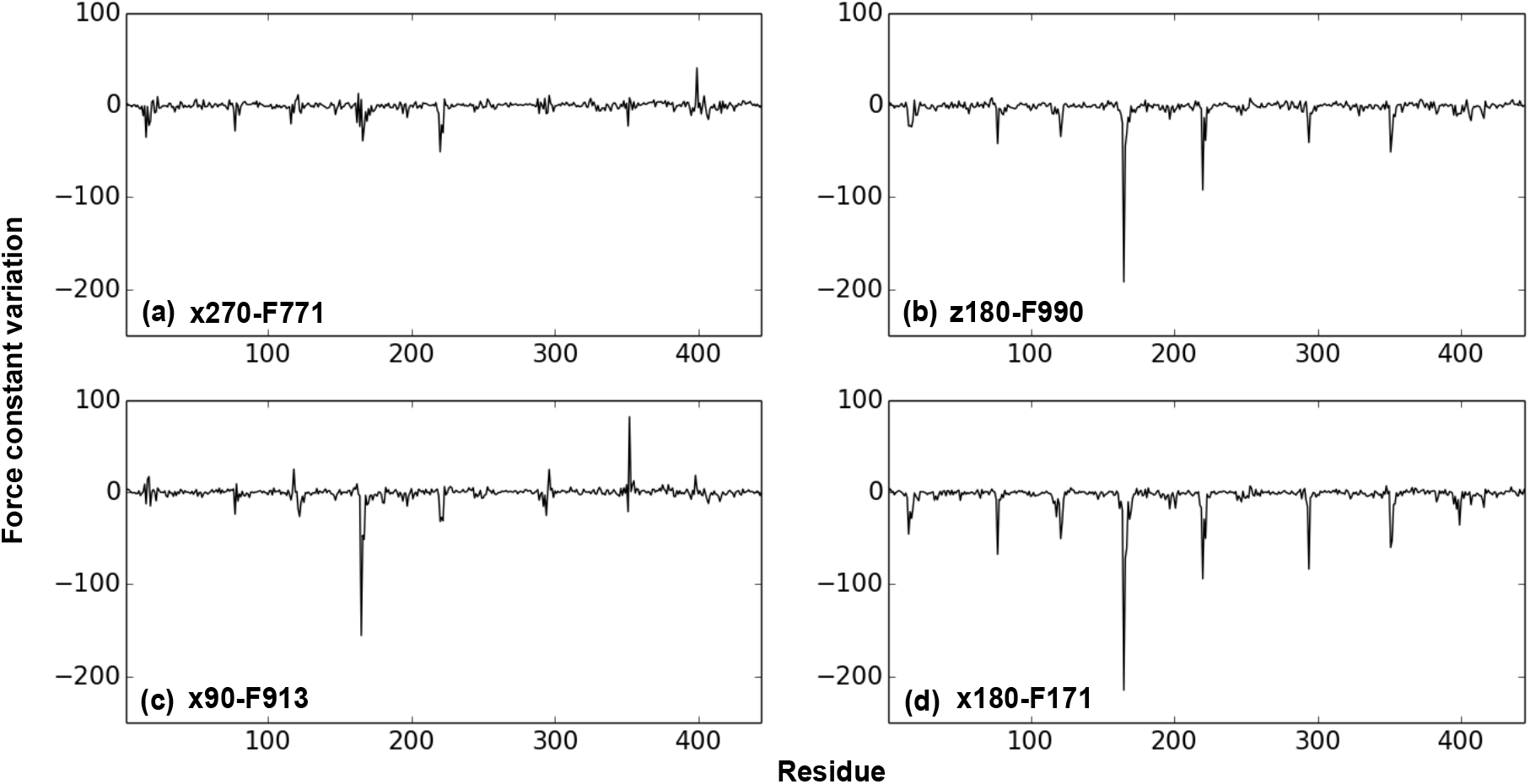
Force constant variations (in kcal.mol^−1^.Å^−2^) in four βGA representative structures from the confined trajectories compared to the reference bulk structure. Trajectories: (a) x270 (b) z180 (c) x900 (d) x180.

The mechanical changes in βGA can also be related to its adsorption behavior during the MD simulations. For example, during trajectory x270, βGA will detach itself from the SAM functionnalized surface after 50 ns, and will spend the rest of the simulation without forming new contacts with either surface. In Figure 7a, we can see how the selected structure (corresponding to frame 771, that is for 77 ns of simulation) only displays minor mechanical variations. On the other hand, structures corresponding to the adsorbed enzyme, either on the SAM or on the bare gold surface, present large mechanical changes. Interestingly, it seems that the mechanical impact of adsorption might also be linked to the adsorbing surface. Frames z180F990, x90F913 and x180F171 correspond to structures of βGA adsorbed on the bare gold surface, and they also present the most important rigidity loss for catalytic residue Glu166 (see Figures 7b-d). This flexibilization of the catalytic center might be detrimental to the enzyme function.

Depending on the protein position between the two solid surfaces (see Figure SI-3 for a global view), the 13 representative structures extracted from the confined trajectories can be classified as unbound (structures z0F118, x90F265, x180F62 and x270F771), SAM-bound (z0F118, z0F461, z180F19, z180F186 and x0F270) or gold-bound (z180F990, x90F913, x180F171 and x180F797). While the average force constant in the unbound and SAM-bound structures remains very close to what was observed in the reference structure from the bulk simulations (with values of 25.5, 25.6 and 25.7 kcal.mol^−1^.Å^−2^ respectively), the gold-bound structures present a reduced average force constant of 23.5 kcal.mol^−1^.Å^−2^. Interestingly, this increase in the protein flexibility when adsorbed on a bare gold surface compared to the SAM-covered surface was also observed in an earlier modeling study on cytochrome *c*,^63^ and again, this perturbation of the enzyme mechanics is likely to impact its catalytic activity.

## 4. Conclusions

Understanding the interaction of enzymes with solid surfaces is a central issue in the biomaterials field. Here, we developed a multi-scale approach, based on all-atom Molecular Dynamics and coarse-grain Brownian Dynamics simulations, to investigate the binding of β-Glucosidase A on two solid surfaces: a bare gold surface and a SAM-OH functionalized gold surface. We performed six 100 ns simulations using different starting orientations for βGA. While these trajectories are too short to establish a representative sampling of the confined enzyme orientational and conformational landscape, as they only describe the early stage of adsorption, their analysis enable us to highlight a different behavior for βGA on the SAM-OH and on the bare gold surfaces. All the MD trajectories start with the enzyme located near the SAM-OH surface, the protein will desorb from the SAM in four cases out of six, and in three cases it will permanently bind to the bare gold surface. Looking at the protein/surface interaction energies, we can see that the interaction of βGA with the bare gold surface is indeed much stronger (by a factor of 4) than with the SAM-OH surface. But this increase in stability goes with a loss of specificity in the bound-enzyme orientation, as the binding spots for βGA on bare gold are distributed all over the protein surface. While adsorption on both surfaces seems to have little impact on the protein conformation, our coarse-grain study of the enzyme shows that it is nonetheless sufficient to induce noticeable changes into its mechanical properties. In particular, adsorption on the bare gold surface leads to an important decrease in the rigidity of catalytic site residues, which might in turn lead to a reduced enzyme activity. One should also note that in the case of βGA, the broad size of the catalytic site,^48, 87^ which is located at the entrance of the central β-barrel, and its structural stability ensure that it remains open at all times. However, many proteins comprise a buried active site that is connected to the surface through internal tunnels and cavities (like globins^88^ or hydrogenases^38^). In that case, one should also specifically investigate how the adsorption process might impact the tunnels width and topology, as a disrupted network is likely to perturb the access to the active site and the enzyme’s catalytic activity.

Altogether, our results show that the efficient immobilization of enzymes on surfaces remains a complex issue. When designing an enzyme-based device, one must find an acceptable trade-off between stability and activity, as was recently illustrated by the experimental work of Weltz et al.^89^ In this perspective, molecular modeling tools have a lot to offer, since they permit to simultaneously retrieve information regarding both the immobilized enzyme structural stability and its function.

## Supporting Information

Surface electrostatic potential of βGA, representative structures used for the coarse-grain mechanical calculations, interaction energy between βGA and the solid surfaces, dipole moment orientation along time, βGA radius of gyration, technical note on the gold force-field parameters choice.

## Acknowledgments

This work was supported by the ANR (ENZYMOR-ANR-16-CE05-0024) and by the “Initiative d’Excellence” program from the French State (Grant “DYNAMO”, ANR-11-LABX-0011-01). Simulations were performed using the HPC resources from LBT/HPC.

