## Supplementary Information for "Between two walls: Modeling the adsorption behavior of β-glucosidase A on bare and SAM-functionalised gold surfaces"

**Figure SI-1:** Surface electrostatic potential resulting from APBS calculations on two opposite faces of βGA . Positive potentials are shown in blue and negative ones in red.

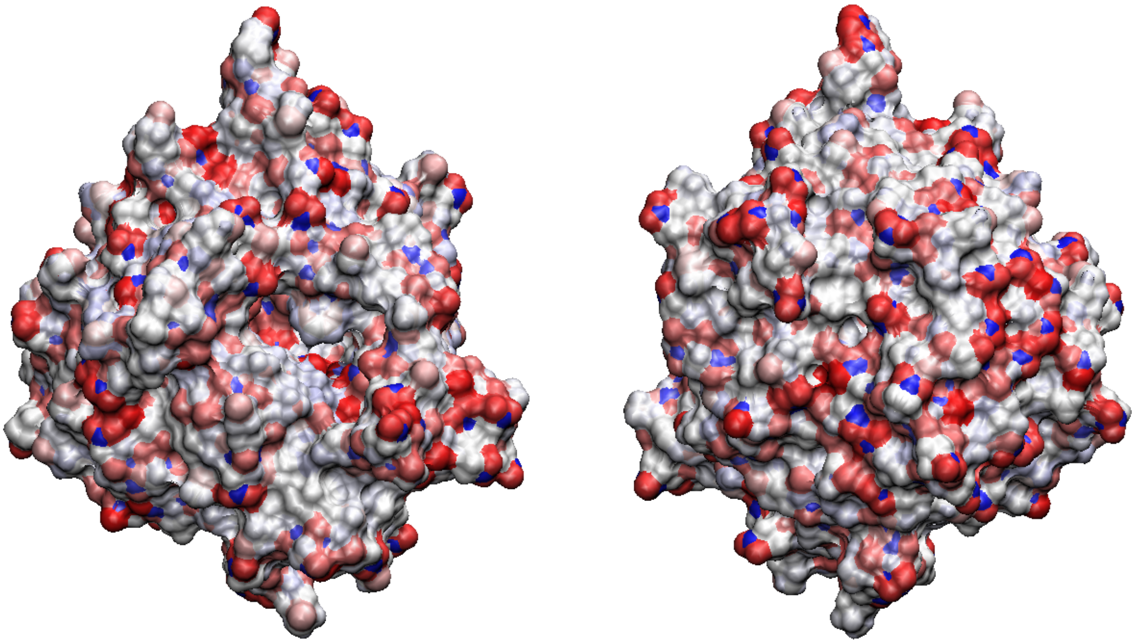

**Figure SI-2:** Cartoon representations of the starting position of βGA in the six confined trajectories.

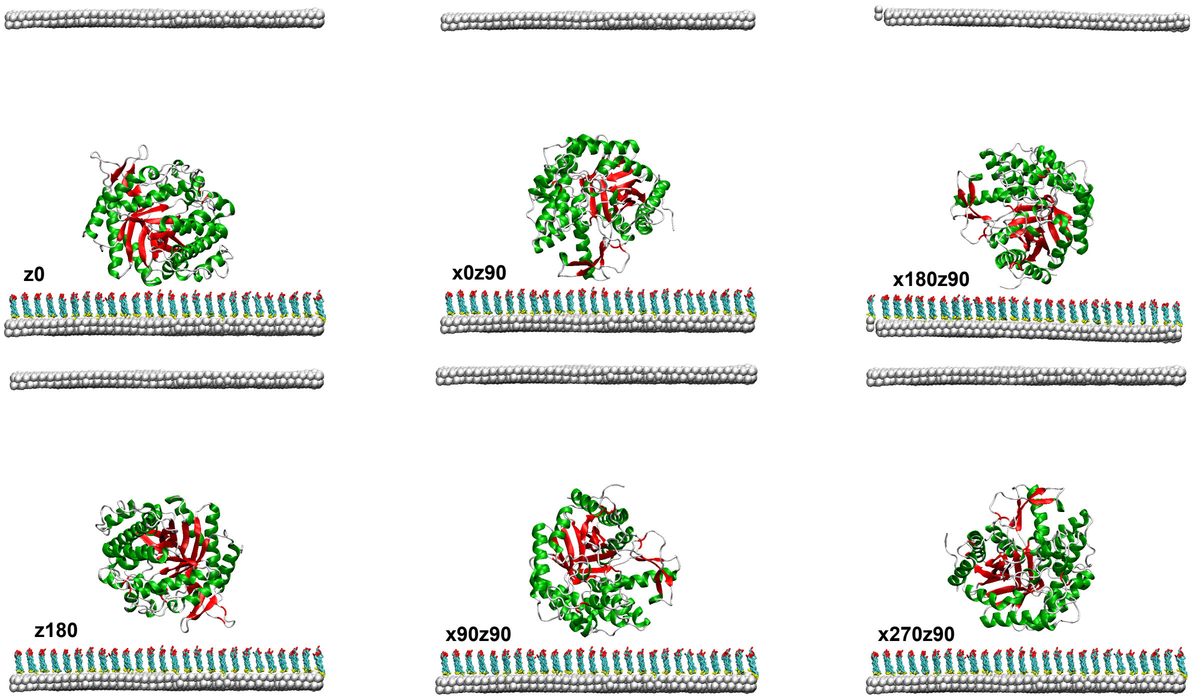

**Figure SI-3:** Cartoon representations for the 13 representative structures of βGA used to investigate the enzyme's mechanical properties.

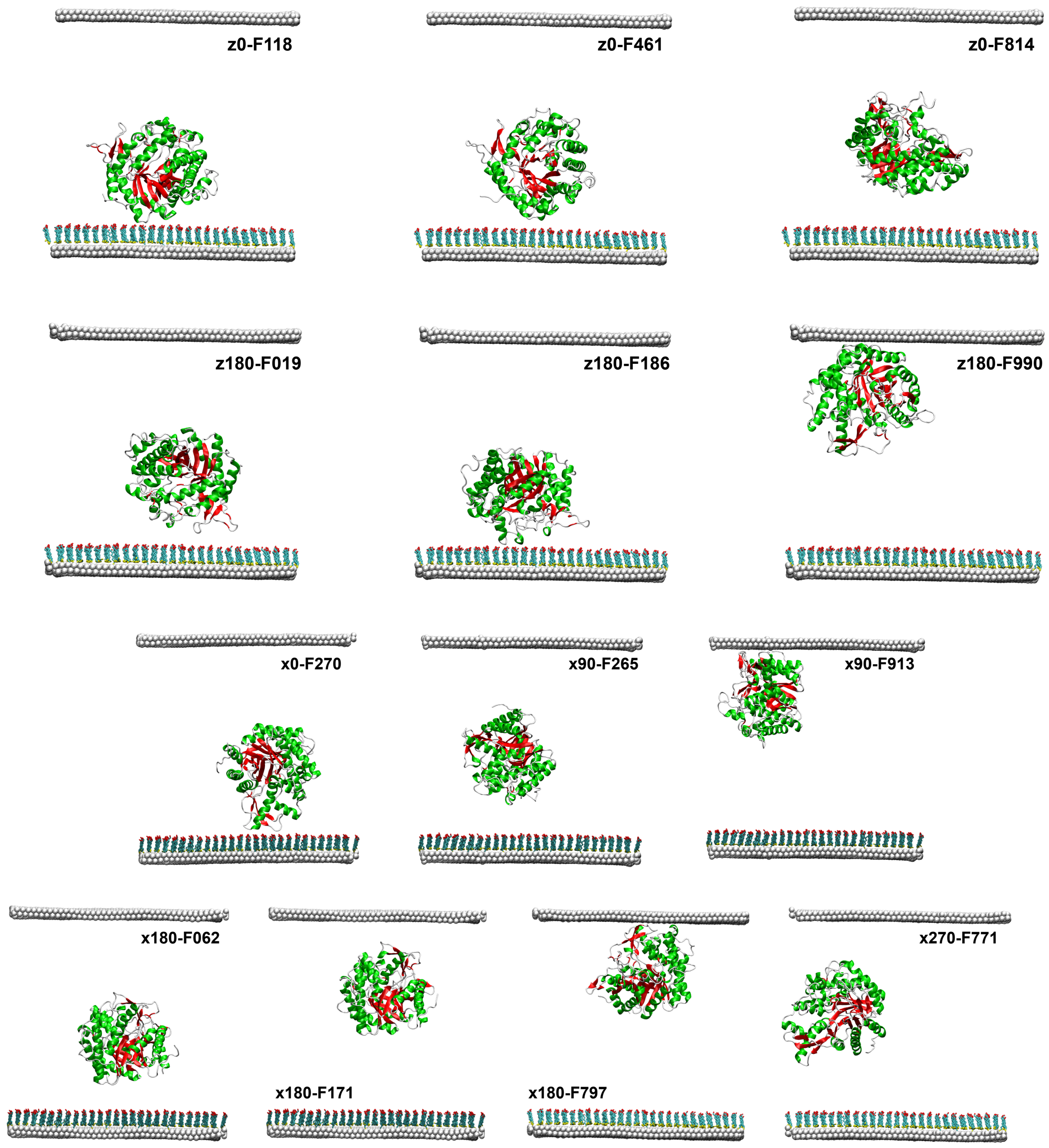

**Figure SI-4:** Black lines (left vertical axis): Distance between the SAM surface and the βGA center of mass as a function of time.

Blue and red lines (right vertical axis): Interaction energy between the enzymes and the SAM-covered surface, or the bare gold surface respectively, as a function of time.

Confined simulations: (a) z0 (b) z180 (c) x0 (d) x90 (e) x180 (f) x270

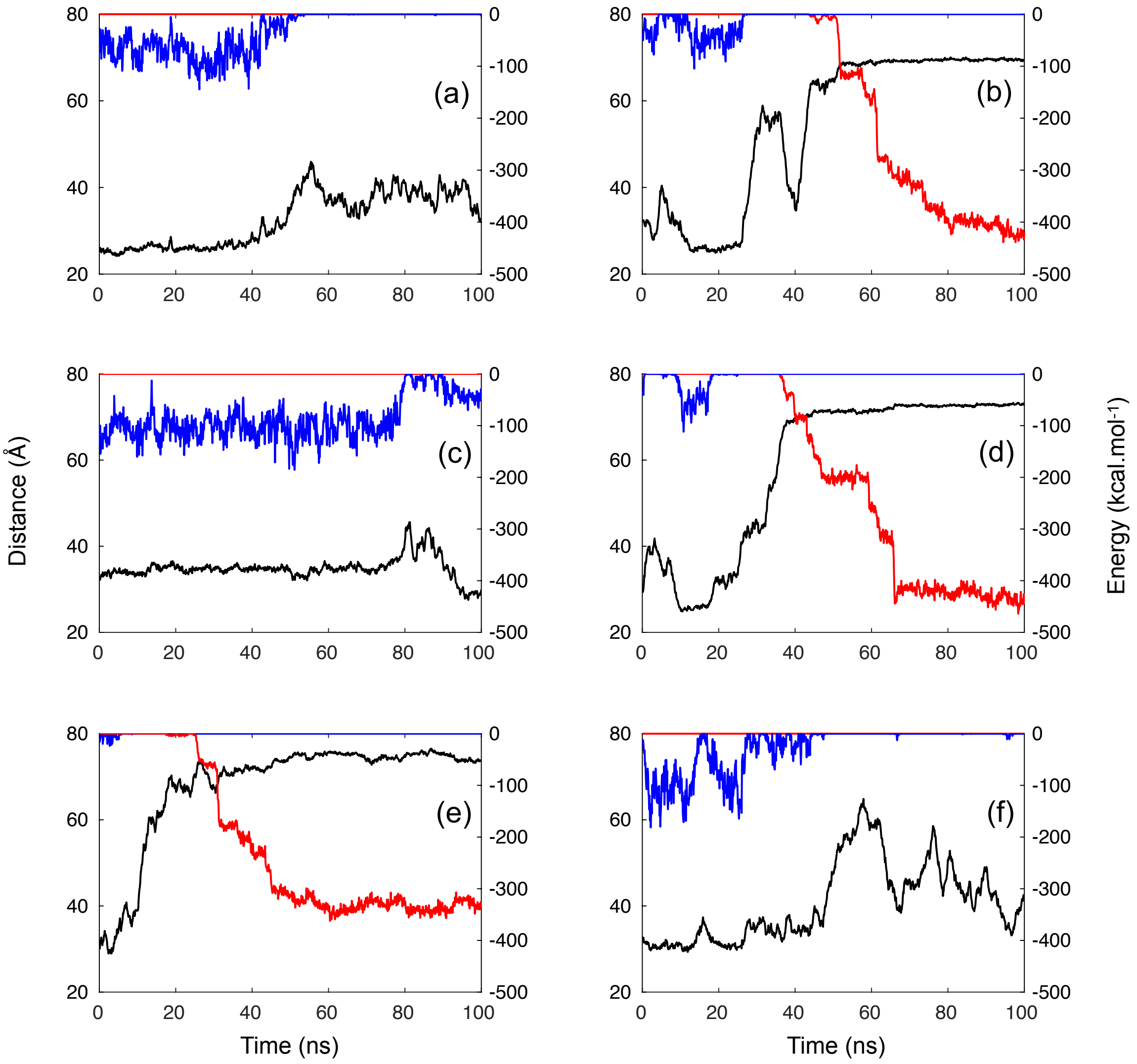

**Figure SI-5:**

(a) Distribution of the βGA dipole moment orientation with regard to the protein structure during the bulk MD trajectory.

(b) Distribution of the βGA dipole moment orientation with regard to the protein structure during the six confined MD trajectories.

(c) Angle between the dipole moment and the Ile119-Asn353 vector as a function of time for all MD simulations.

(d) Definition of the tilt ($\theta$) and twist($\varphi$) angles that were used to monitor the βGA orientation with regard to the SAM-covered surface during the simulations. The enzyme's dipole moment is shown as a blue arrow, and the Cα atoms of residues Ile119 and Asn353 as blue van der Waals spheres.

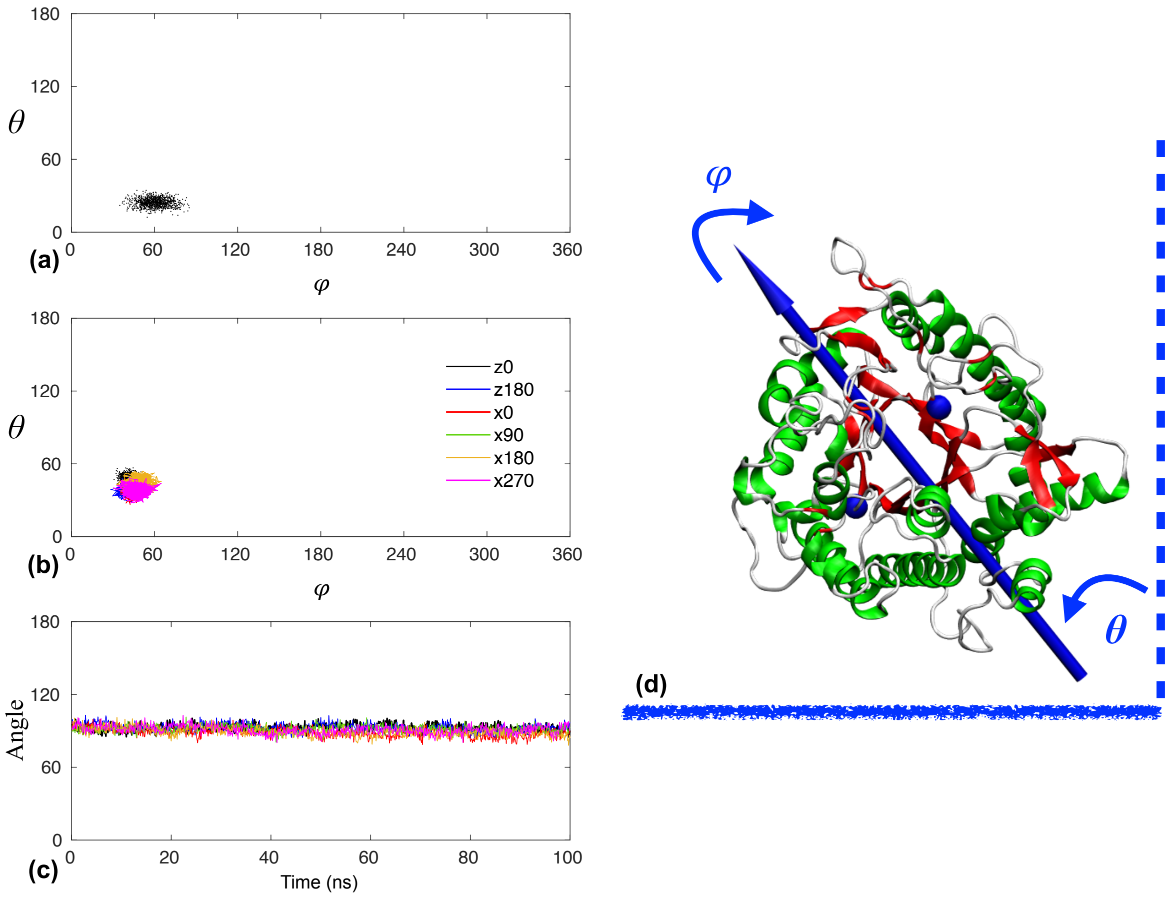

**Figure SI-6:** Average residues contact frequencies with the solid surfaces. (a) Contact frequencies with the SAM-covered gold surface, average values over all six confined trajectories. (b) Contact frequencies with the bare gold surfaces, average values over trajectories z180, x90 and x180.

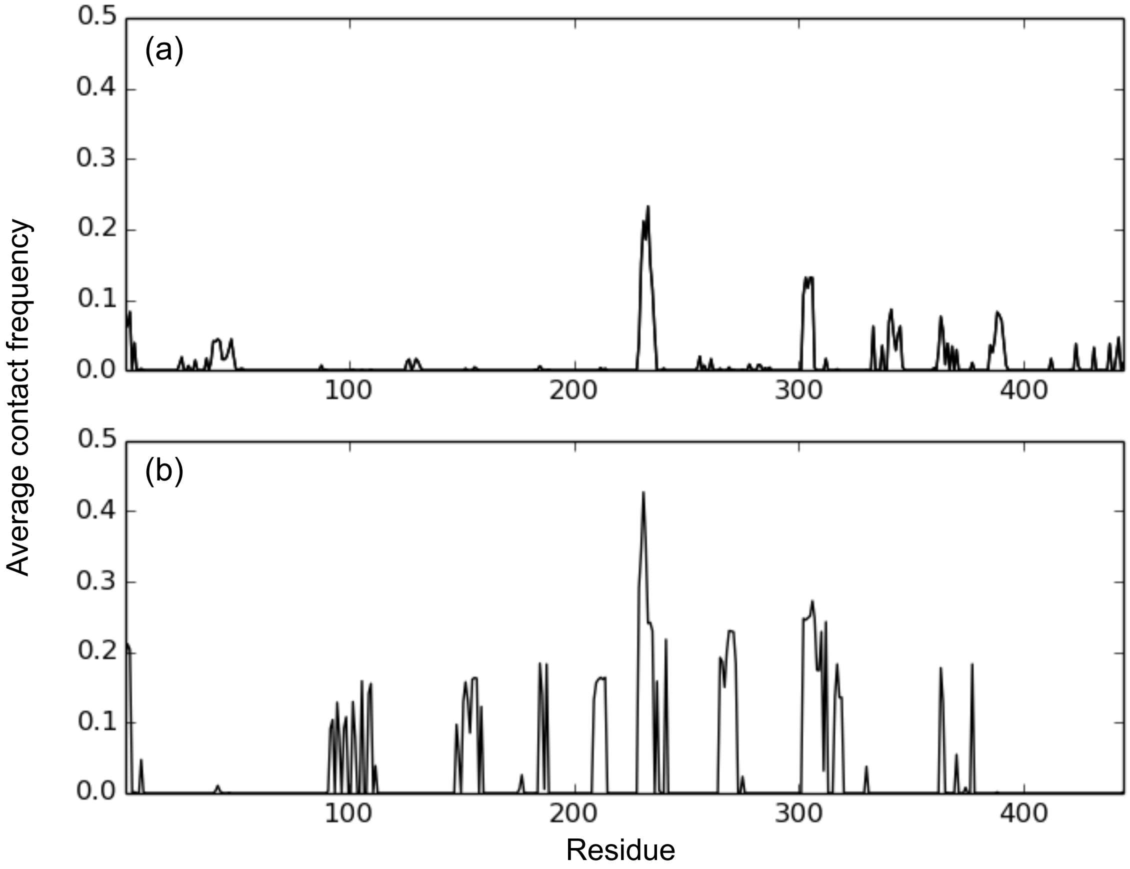

**Figure SI-7:** Density distributions for the βGA radius of gyration. (a) bulk trajectory, (b) confined trajectories.

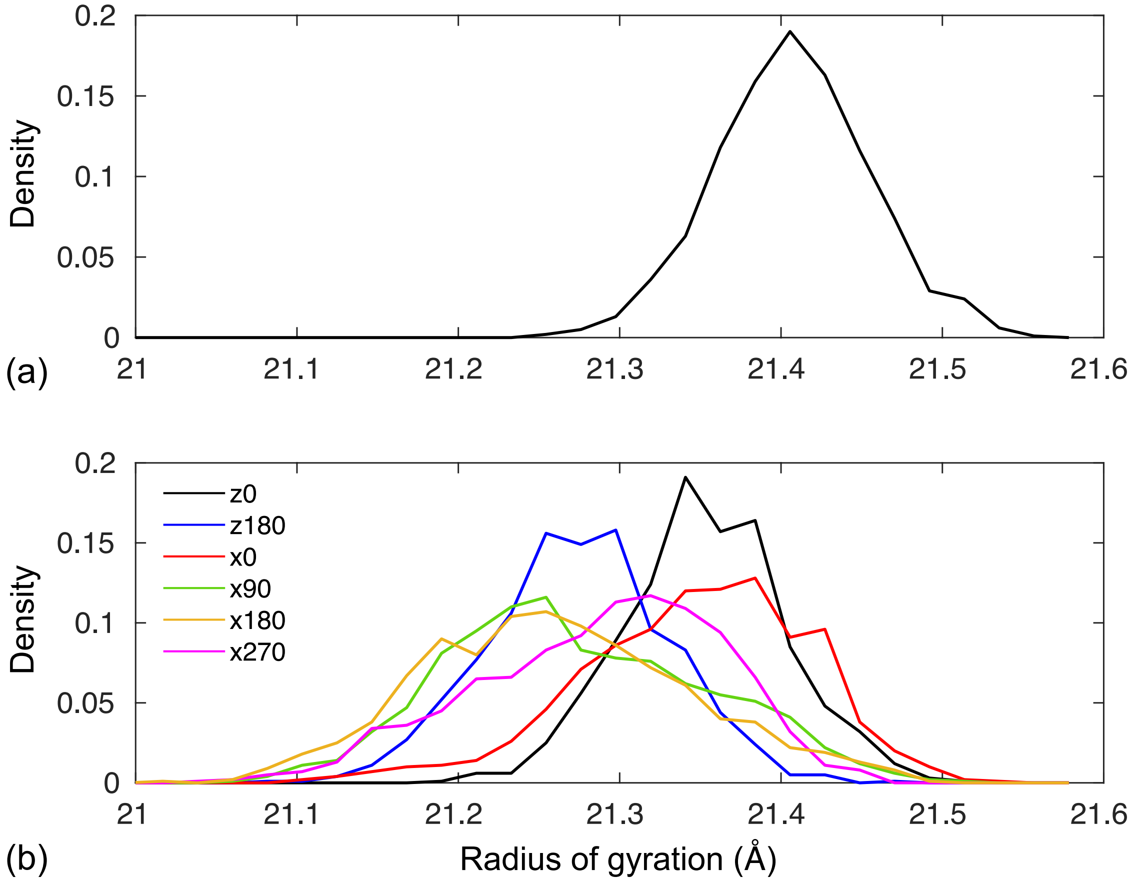

**Figure SI-8:** Force constant variations (in kcal.mol$^{-1}$.Å$^{-2}$) in the 13 βGA representative structures from the confined trajectories compared to the reference bulk structure.

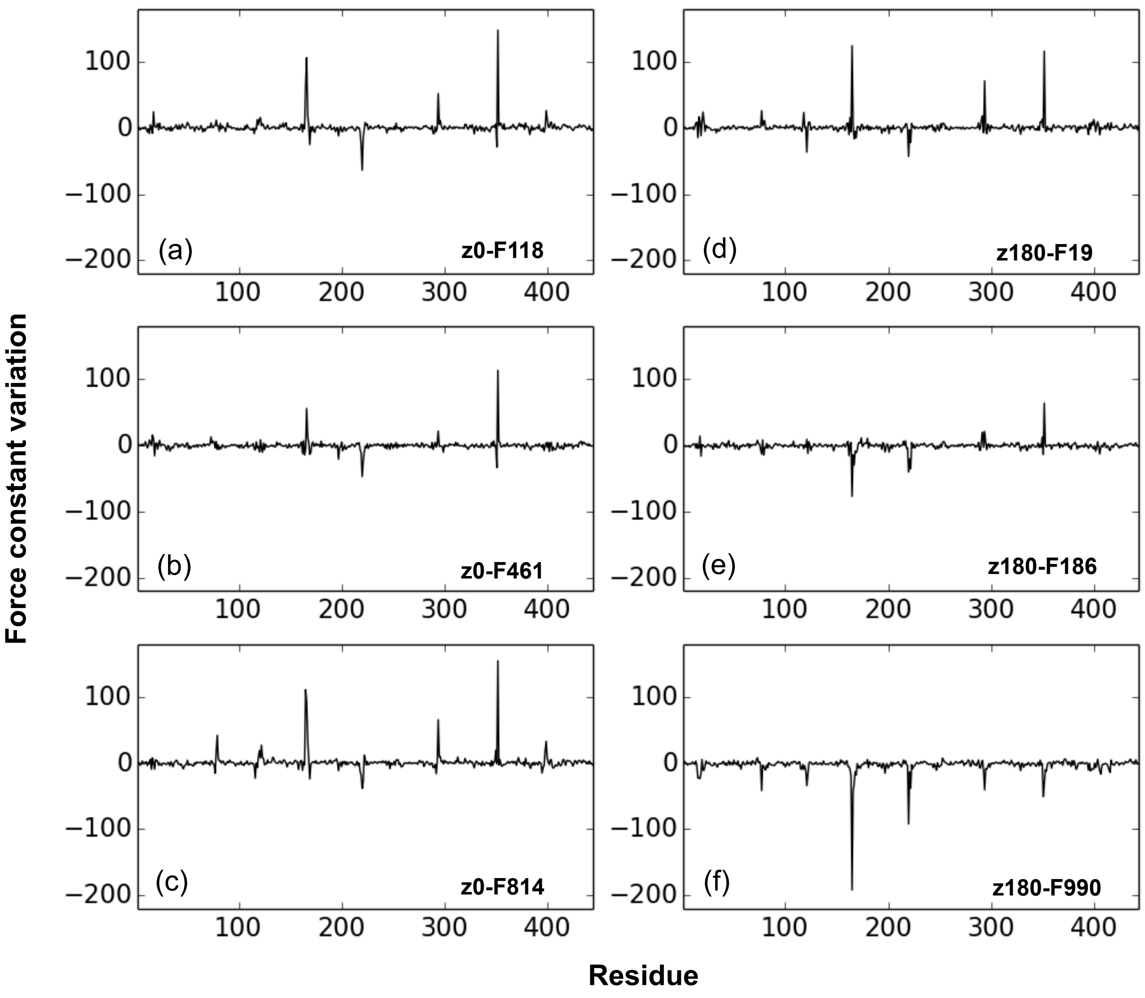

**Figure SI-8 (continued):** Force constant variations (in kcal.mol$^{-1}$.Å$^{-2}$) in the 13 βGA representative structures from the confined trajectories compared to the reference bulk structure.

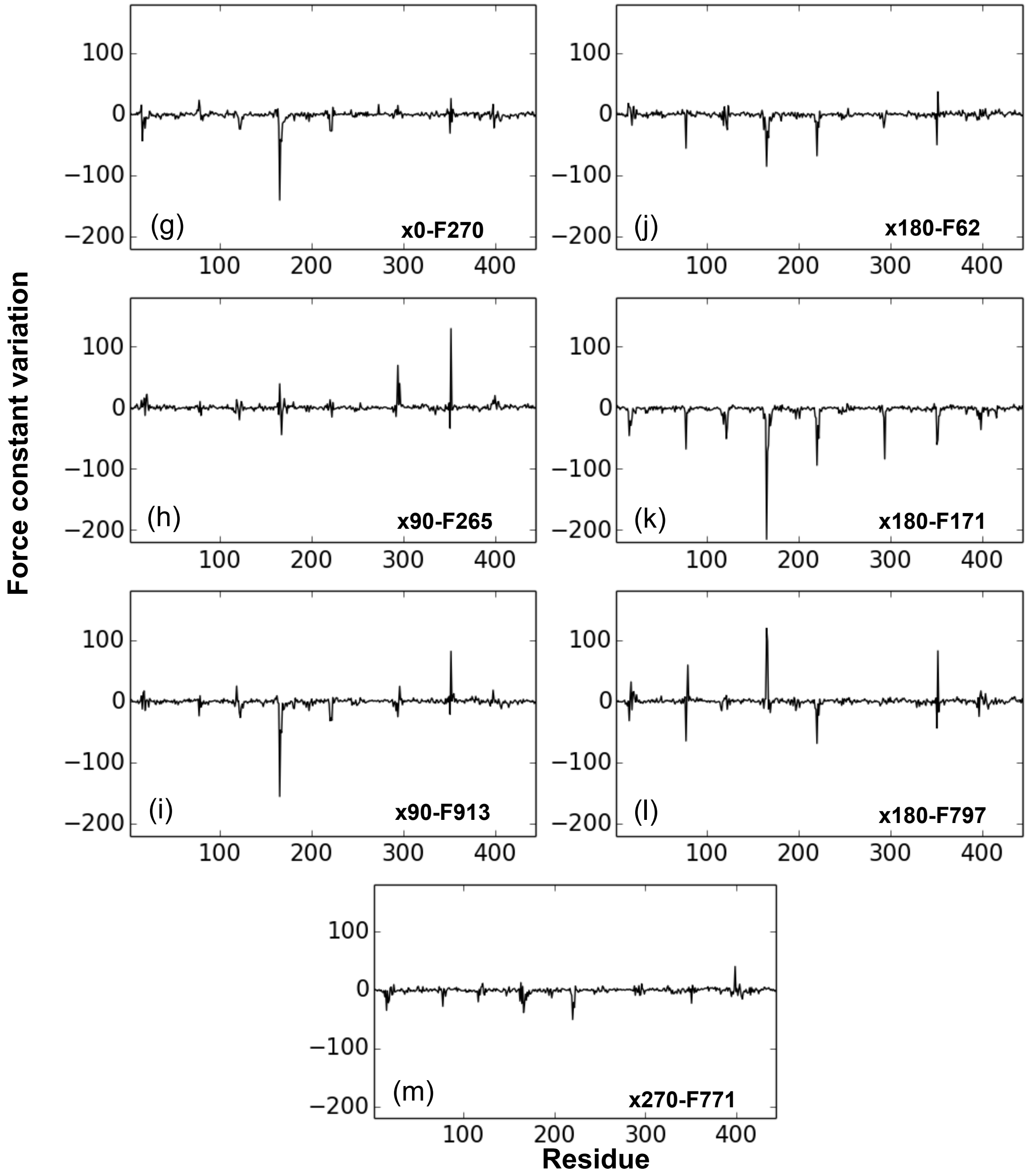

**Technical note on the gold force-field parameters choice:**

During Molecular Dynamics simulations, the potential energy of the system is computed through a dedicated force-field. The choice of gold parameters needs a peculiar attention, specifically for its Lennard-Jones terms where several sets of parameters are available and summarized in table 1. These parameters have their own coherence so that it has no sense to mix their data.

*Table 1: Available Lennard-Jones parameters and their references for gold atoms on Au{111} surface (first 3 columns) and Au{001} (last column).*

| Term/Ref | Agrawal[1] | Interface FF[2,3] | Golp[4] | Vila Verde[5] |
|---|---|---|---|---|
| $\sigma_{Au}$ | 2.569 Å | 2.951 Å | 1.796 Å | 3.293 Å |
| $\varepsilon_{Au}$ | 0.458 eV | 0.229 eV | 0.007 eV | 0.046 eV |

Treatment of electrostatics, including electronic polarization, is one of the most important requirements for an accurate modeling of biomolecules. However, even if much efforts have been done during several years to develop polarizable force-fields, their computational cost is still important. Besides, a previous study[6] mitigates the influence of polarizability during the adsorption process of molecules on an Au{111} surface in a solvated system since the interfacial water molecules drastically abate this effect. For these reasons, it was decided not to use the polarizable force-field Golp[4,7].

To choose between the three other possibility (Agrawal, Interface FF and Vila Verde), we took into account experimental background on this topic[8] and our previous computations[9,10] with plane-wave periodic density functional theory (DFT). From these, we considered a gold Au{111} plane with 2 slab layers functionalized with 1-mercapto-6-hexanol molecules. These last take an asymmetric bridge position on the surface providing a S−Au bond length of 2.392 Å and a tilt angle of 49.6° between the main axis of the molecule and the surface normal vector. According to these data, we chose the parameters of Agrawal because the sigma value is much close to our Au-S interaction which occurs in our 1-mercapto-6-hexanol functionalized gold surface. This choice was also set in other works on nucleic-acids, proteins, polymers and organic molecules (for instance see[11–19]) but other sets of parameters also provided meaningful results in gold/protein adsorption studies (for example see[3,20,21]).

Finally, a RMSD of 1.03 Å was recorded between the whole gold functionalized surface issue from the propagation of the DFT calculation and the optimized structure without any restraint. This was found adequate to pursue the molecular dynamics calculations.
